# Synchronised spiking activity underlies phase amplitude coupling in the subthalamic nucleus of Parkinson’s disease patients

**DOI:** 10.1101/433334

**Authors:** Anders Christian Meidahl, Christian K.E. Moll, Bernadette van Wijk, Alessandro Gulberti, Gerd Tinkhauser, Manfred Westphal, Andreas K. Engel, Wolfgang Hamel, Peter Brown, Andrew Sharott

## Abstract

Both phase-amplitude coupling (PAC) and beta-bursts in the subthalamic nucleus have been significantly linked to symptom severity in Parkinson’s disease (PD) in humans and emerged independently as competing biomarkers for closed-loop deep brain stimulation (DBS). However, the underlying nature of subthalamic PAC is poorly understood and its relationship with transient beta burst-events has not been investigated. To address this, we studied macro- and micro electrode recordings of local field potentials (LFPs) and single unit activity from 15 hemispheres in 10 PD patients undergoing DBS surgery. PAC between beta phase and high frequency oscillation (HFO) amplitude was compared to single unit firing rates, spike triggered averages, power spectral densities and phase-spike locking, and was studied in periods of beta-bursting. We found a significant synchronisation of spiking to HFOs and correlation of mean firing rates with HFO-amplitude when the latter was coupled to beta phase (i.e. in the presence of PAC). In the presence of PAC, single unit power spectra displayed peaks in the beta and HFO frequency range and the HFO frequency was correlated with that in the LFP. Finally, PAC significantly increased with beta burst-duration. Our findings offer new insight in the pathology of Parkinson’s disease by providing evidence that subthalamic PAC reflects the locking of spiking activity to network beta oscillations and that this coupling progressively increases with beta-burst duration. These findings suggest that beta-bursts capture periods of increased subthalamic input/output synchronisation in the beta frequency range and have important implications for therapeutic closed-loop DBS.

**Significance statement:** Identifying biomarkers for closed-loop deep brain stimulation (DBS) has become an increasingly important issue in Parkinson’s Disease (PD) research. Two such biomarkers, phase–amplitude coupling (PAC) and beta-bursts, recorded from the implanted electrodes in subthalamic nucleus in PD patients, correlate with motor impairment. However, the physiological basis of PAC, and it relationship to beta bursts, is unclear. We provide multiple lines of evidence that PAC in the human STN reflects the locking of spiking activity to network beta oscillations and that this coupling progressively increases with the duration of beta-bursts. This suggests that beta-bursts capture increased subthalamic input/output synchronisation and provides new insights in PD pathology with direct implications for closed-loop DBS therapy strategies.

## Introduction

Deep brain stimulation (DBS) is a well-established treatment of severe Parkinson’s disease (PD). In recent years several putative biomarkers have emerged for closed-loop applications and new experimental interventions that might further improve DBS efficacy. Beta (13-35 Hz) power in local field potential (LFP) recordings from the subthalamic nucleus (STN) of PD patients correlates with akinetic/rigid motor impairment and is attenuated by DBS and levodopa medication (Kühn et al., 2006, 2008, 2009; Ray et al., 2008; Steiner et al., 2017). Changes in power of high frequency oscillations (HFOs, 100-300 Hz) have been observed between on and off dopaminergic treatment (Lopez-Azcarate et al., 2010) and during movement initiation and planning (Combrisson et al., 2017). Additionally, neuronal firing patterns in the STN have been linked to symptom severity in PD with intra-burst rates and beta-oscillating unit activity positively relating to bradykinesia, rigidity and axial scores (Sharott et al., 2014).

Phase-amplitude coupling (PAC), where the phase of a low frequency is coupled to the amplitude of higher frequency oscillation, is believed to serve a role in a variety of physiological brain functions (Cohen et al., 2009; Colgin et al., 2009; Tort et al., 2009; Carracedo et al., 2013; Richardson et al., 2017). In PD, however, beta-HFO PAC in the STN (Lopez-Azcarate et al., 2010; van Wijk et al., 2016) and beta-gamma PAC in motor cortex (Shimamoto et al., 2013; de Hemptinne et al., 2015) have been shown to correlate with motor symptom severity in patients and to develop in non-human primates in the STN after dopamine depletion (Escobar et al., 2017). PAC in the STN is attenuated by dopamine replacement (Lopez-Azcarate et al., 2010; van Wijk et al., 2016), and PAC in the motor cortex is reduced by STN DBS (de Hemptinne et al., 2015), providing further evidence for PAC as a biomarker of PD. Whether these correlations reflect a mechanistic relationship with PD symptoms, and how PAC relates to other established biomarkers of PD symptoms, has not been fully determined.

Interpreting the importance of PAC to PD pathophysiology is complicated by conflicting explanations for the underlying mechanism. Low frequency phase in PAC is often thought to reflect the synchronisation of synaptic potentials that can drive coordinated firing of local neurons (Canolty and Knight, 2010). This mechanism is likely to underlie the phase of beta oscillations in STN LFPs in PD, which have fixed timing relationships to firing of single neurons (Steigerwald *et al.*, 2008; Shimamoto *et al.*, 2013; Yang *et al.*, 2014; Lipski *et al.*, 2017; Sharott *et al.*, 2018). However, recent studies have emphasised the transient nature of beta oscillations, which occur in “bursts” of high amplitude (Tinkhauser *et al.*, 2017a; Tinkhauser *et al.*, 2017b). It is currently unclear as to whether PAC is restricted to such bursts or can occur independently of the amplitude of the beta oscillation.

Investigations into the relationship between HFOs and spiking activity in the STN have produced conflicting results. Wang and colleagues (Wang et al., 2014) found that HFOs were independent spatiotemporal phenomena from neuronal spiking. These authors concluded that large pools of desynchronised neuronal clusters give rise to nonstationary HFOs in the LFP that are distinct from the locally-synchronised multiunit activities and are also phase locked to the beta oscillation (Weinberger et al., 2006; Sharott et al., 2014). However, a recent computational study found that synchronized single-cell bursting led to PAC that closely resembled that of parkinsonian mice and primates and could produce PAC resembling that seen in patients (Sanders, 2016).

Using intraoperative macro- and microelectrode recordings in PD patients off medication, we demonstrate that HFO amplitude is related to the firing rate and pattern of single STN neurons in the presence, but not absence of PAC. In addition, PAC and HFO amplitude are higher during beta bursts, suggesting that PAC occurs during enhanced beta synchronisation. These findings suggest that STN PAC could be a proxy for the locking of neurons to network oscillations.

## Methods

Local field potentials and single unit recordings were captured simultaneously from subthalamic nuclei in 15 hemispheres in 10 patients during surgical DBS electrode placement. All patients (6 female, 4 male; mean ± SD age: 66.8 ± 3.4 years) suffered from advanced idiopathic PD and were withdrawn from anti-parkinsonian medication the night before surgery. For patient details see Supplemental Table 1. The surgical procedure yielded 231 local field potential recordings with 157 single units extracted from microelectrode recordings (14 hemispheres in 9 patients were used for analysis of single units). All LFPs/spike train pairs analysed in this paper were recorded by a distance of at least 2 mm to remove spiking contaminations of recorded LFPs in the analysis.

### Surgical procedure and microrecordings

Electrophysiological data from all patients in this study have been reported previously (Sharott *et al.*, 2014; Sharott *et al.*, 2018). Operations were performed under local anaesthesia. Details concerning the surgical procedure are reported elsewhere (Hamel *et al.*, 2003; Moll *et al.*, 2014; Sharott *et al.*, 2014; Sharott *et al.*, 2018). Dopamine agonist treatment was stopped more than 7 days before the operation. 10-15 minutes prior to the start of microelectrode recordings at the level of the thalamus (usually 6-12 mm above the centre of the STN as delineated on MRI), systemic sedation with low dose remifentanyl (0.01 - 0.1 μg/kg/min) was completely stopped. No sedatives or anesthetics were administered during the microelectrode mapping procedure. Participation in the study extended the surgical procedure by approximately 15-25 minutes.

Microelectrode recordings were performed along five parallel tracks arranged in a concentric array (MicroGuide, Alpha-Omega, Nazareth, Israel) (Supp. Fig. 1). Four outer platinum-iridium electrodes (impedance = 0.7 ± 0.2 (range, 0.2 - 1.25) megaOhm at 1000 Hz; FHC Inc., Bowdoinham, ME, USA) were separated by 2 mm from a central one, which was aimed at the STN surgical target. Signals were amplified (×20.000), bandpass-filtered (0-300 Hz) and digitized (sampling rate: 24 kHz). An electrode was identified as having entered the STN following a clear increase in the size of the background activity (Moran *et al.*, 2006). STN units were distinguished by their tonic irregular, oscillatory or bursty discharge pattern (Supp, Fig. 1), and this was clearly different from slower bursting and single-spiking of neurons in the overlying thalamus and zona incerta, and from the regular, high frequency spiking of more ventral substantia nigra pars reticulata neurons.

## Data selection and processing

All data were collected as part of our routine neuronavigation process which aims to precisely identify the STN. Recordings were only included if they were made during sustained periods during which patients were awake and highly co-operative. In total, 157 STN unit recordings were used in this study. In some recordings, patients were engaged in a simple, brief movement task (flexion/extension of wrist or ankle) as part of the routine mapping procedure. Such epochs comprised a relatively small proportion of individual recordings and we have demonstrated previously that beta oscillations are present in these recordings (Sharott *et al.*, 2014; Sharott *et al.*, 2018). Spike detection was performed offline using a voltage threshold method (Offline-Sorter, Plexon Inc., Dallas, TX, USA). The threshold value was set sufficiently high relative to the noise level to avoid false-positives (threshold values > 4 SD of the background level were used). When possible, single unit activities (SUAs) were then separated by manual cluster selection in 3D feature space on the basis of several parameters including signal energy, principal components, peak time and the presence of a central trough in the autocorrelogram. Over half of the unit activities were characterized as SUAs (n = 157). Any portions of injury discharge were discarded.

### Phase amplitude coupling and neuronal activity

All computation was done in Matlab (R2017a, The Mathworks Inc., Natick, USA). In analyses comparing measures of neuronal activity to PAC, PAC was computed using the General linear model (GLM) method (van Wijk et al., 2015) to allow parametric estimation of significance. After removing 50 Hz harmonics using a 4^th^ order Butterworth filter the LFP signals were initially bandpass filtered around centre frequencies between 10-35 Hz with 1 Hz increments to obtain the beta component and between 100-300 Hz with 4 Hz increments to extract the HFO component. Filter bandwidths were set at ± 1 Hz and ± 35 Hz, respectively, and the instantaneous phase was extracted for the beta components via and amplitude of HFO components via

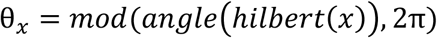

and amplitude of HFO components via

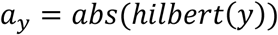

with *x* representing the beta, and *y* the HFO signal components (see Supplementary figure 1). The general linear model:

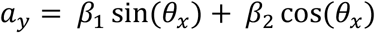

was then applied to all phase and amplitude frequency combinations. Each frequency combination would then yield a single PAC value from

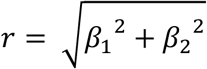

with beta coefficients calculated via least squares.

sin(*θ*_*x*_), cos(*θ*_*x*_) and *a*_*y*_ were all transformed to have zero mean and unit variance, for example: *a*_*y*_ = (*a*_*y*_ − mean(*a*_*y*_))/std(*a*_*y*_). To test for significance the time series was divided into non-overlapping epochs of 3 second durations to allow for parametric statistical testing, yielding separate GLMs and regression coefficients across epochs. The recordings had their edges (2*sampling rate) removed to eliminate filter artefacts. An F-test was applied to test for beta-coefficient consistency across epochs. PAC was defined to be significantly present if a cluster of at least 30 contiguous significant frequency bins of p-values < 0.01 was observed. Data that fell below 10 connecting significant frequency bins of p-values < 0.01 were categorised as non-significant (see figure 1). Data that fell in between were classified as intermediate and excluded from further analysis. Based on this significance testing, the recordings were allocated to groups of “significant” or “non-significant PAC”.

**Figure 1.**
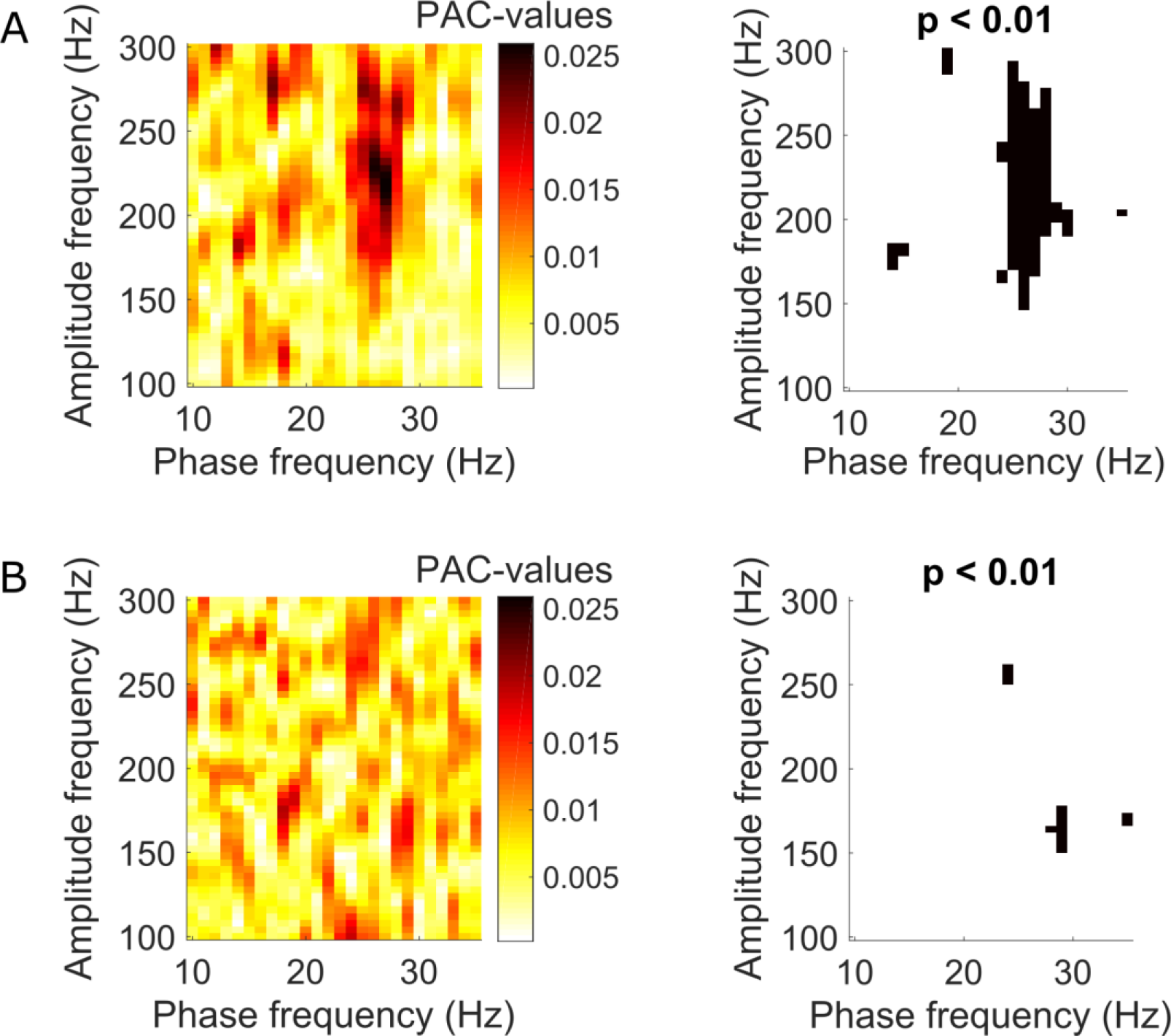
Examples of significant and non-significant PAC-clustering. **A)** Example PAC comodulogram from one recording in one patient with significant coupling between the phase of beta frequencies and HFO amplitude. Frequency bins indicated in black were significant for p < 0.01. **B)** Example comodulogram in the same patient from a different recording without significant PAC-clustering.

### Spike-amplitude coupling

The GLM was further extended by including spiking as an additional predictor to test for Spike-amplitude coupling:

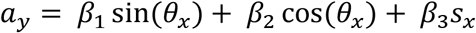

with *s*_*x*_ representing the Z-scored spiking time series. Spike-amplitude coupling was then represented by Spike_AMP_ = *β*_3_ with values ranging from −1 to 1. For individual recordings, a t-test was performed to test for *β*_3_ consistency across epochs for each HFO frequency, to determine levels of significance. For the spike-amplitude-coupling group figures, all the spike-amplitude plots were averaged for the significant and non-significant PAC groups respectively, and a two-sample t-test was performed to determine levels of spike-amplitude significance.

### Spike-triggered averages (STAs)

STAs were computed by triggering the amplitude envelope of the band-pass filtered (4th-order Butterworth) HFO (100-300 Hz) at the time of each action potential from a single unit, where the unit and LFP signals were separated by at least 2 mm. The HFO amplitude was averaged around each single-unit spike with time zero marking spike onset. For averaged group data, a two-sample t-test was performed at spike onset.

### Comparison of firing rates and HFO amplitude within recordings

Mean firing rates and HFO amplitudes were calculated from single unit and LFP recordings in non-overlapping 3 second epochs. The HFO 180-220 Hz amplitude was chosen for comparison as this was the frequency range where the highest degree of significant PAC was found overall. HFO amplitude was extracted as described in the GLM-method section and averaged for corresponding epochs.

*Power spectral densities* of single units were calculated by use of the Welch periodogram method (Matlab function pwelch.m function, using a Hanning window with 50% overlap, window size = 0.2*sampling frequency).

### Spike-phase locking

The Hilbert transform was used to extract phase from band-pass filtered LFP signals (beta-frequency corresponding to highest PAC value +/− 5 Hz) and the phases at the times of each action potential were extracted to construct phase histograms. A Rayleigh’s test for non-uniformity was applied to determine if the spike-phase locking was significant using the Circstat Matlab toolbox (Berens, 2009). Phase histograms were also made from the Hilbert extracted beta phase with regards to HFO, which was separated into 30 amplitude bins and plotted against the beta phase cycle.

### PAC during beta bursts

Beta-bursts were identified in accordance with the method used by Tinkhauser and colleagues (Tinkhauser et al., 2017). The rectified LFP signal was filtered around the individual beta peak frequency as identified by wavelet frequency decomposition. The Hilbert amplitude was extracted from this filtered signal and used to identify periods of beta-bursts above the set threshold and duration criteria. As standard criteria burst thresholds were set at 75% with durations above 100 ms (see Supplementary figure 2). Non-bursting periods were identified as periods where the beta Hilbert envelope fell below a 50% threshold. To ensure sufficient data lengths for PAC calculation the bursting and non-bursting periods were concatenated in two different vectors. When comparisons were made between bursting and non-bursting episodes all vector lengths were adjusted to have the same length as the shortest combined vector. The Hilbert phase of the beta component and the phase of Hilbert HFO amplitude were both extracted prior to cutting and concatenating the data to avoid jumps in the signal between periods.

To calculate PAC during bursting and non-bursting periods we chose the phase-locking value (PLV) method between beta phase and the phase of HFO-filtered Hilbert transformed amplitude (Cohen, 2008; Seymour et al., 2017). Like the GLM method, the PLV method is non-susceptible to HFO amplitude levels, which we found to be progressively increased with increased burst thresholds and durations. However, the PLV method has been shown to be superior to the modulation index for short-epoch data (Penny et al., 2008) and the GLM method relies on longer data sets for statistical testing. For the PLV method we tested for statistical significance via 500 runs of surrogate data with circularly shifted permutations of the phase (Penny et al., 2008; Seymour et al., 2017).

The PLV is defined as:

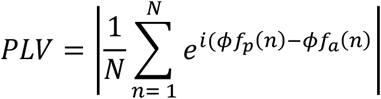

*ϕf*_*a*_ represents the phase time series of the HFO amplitude envelope and *ϕf*_*p*_ the beta phase time series. PAC assumes the value of 1 when the phase series are fully locked and 0 if they are completely desynchronised (Tort et al., 2010; Seymour et al., 2017). Before the *ϕf*_*a*_ by way of the Hilbert transform was extracted from the HFO-amplitude time series, the amplitude signal was high- and lowpass filtered as previously described using Butterworth filters to remove non-beta components (Penny et al., 2008). Absolute PAC values were calculated as the mean of the PAC values in the range of the HFO frequency for the max PAC value ±16 Hz.

For estimation of the mean HFO and beta amplitude during bursting and non-bursting episodes the signals were filtered as previously described for the respective components under the GLM method and, after standardizing the entire time series to have zero mean and unit variance, amplitudes were averaged separately for the bursts and non-bursts periods (Supp. Fig 2).

## Results

### Defining the presence of STN LFP phase amplitude coupling

We analysed 114 micro- and macroelectrode recordings to investigate phase-amplitude coupling of LFP signals in the subthalamic nucleus (Supp. Fig. 1). Recordings were divided based on whether or not significant PAC was detected using significant frequency-bin clustering. Significant PAC-clustering in the STN on a grand average was predominantly found between beta (20-35 Hz) phase and 160-220 Hz amplitude with individual maximum PAC values peaking on average across the HFO frequency range at either 180 or 220 Hz (Fig. 1).

Previous work has suggested that the timing of beta synchronised HFOs and spiking activity in STN are independent (Yang *et al.*, 2014). Our aim was to re-examine this issue and to define precisely how the spiking of STN units is related to PAC. As using spikes and LFPs recordings from the same electrode can obviously introduce contamination between spikes and any other signals, including HFOs, all unit/LFP pairs analysed here were recorded from different electrodes. The configuration of our recording system dictated that these were at least 2mm apart, excluding the possibility of substantial LFP signal contamination (Supp. Fig 1. Einevoll et al., 2013).

### STN firing predicts HFO amplitude only in the presence of PAC

We first examined whether simple correlations in the time domain between HFOs and spike times only occurred in recordings when PAC was present. To achieve this, spike triggered averages of the HFO amplitude (STA-HFO) were compared across PAC and non-PAC recordings (Fig. 2). In single recordings, peaks could be observed at zero lag in the presence of significant PAC (Fig. 2A), but not when PAC was absent (Fig. 2B). Over all recordings, HFO amplitude was significantly locked to spike-onset in the presence of significant PAC, but not in the absence thereof (p = 0.0014, two sample t-test, n(Non-PAC) = 38 n(PAC) = 39, Fig. 2C). In recordings with PAC, secondary peaks could be observed at latencies consistent with the spike-trigged HFO oscillating at beta frequencies (e.g. multiples around ~33 ms corresponding to ~30Hz, Fig 2C, E). Furthermore, the area under the STA-HFO around time zero was positively correlated to the number of connected significant PAC frequency bins (p = 0.0103, r = 0.42811, n = 33, F-test, Fig. 2F), demonstrating that the larger the span of significant PAC across frequencies, the greater the STA HFO amplitude peaks. Importantly, the lack of STA-HFO peaks in the absence of PAC demonstrates that the presence of spikes does not lead to increased HFO amplitude *per se*, rather that spikes and HFOs are specifically associated when PAC is present.

**Figure 2.**
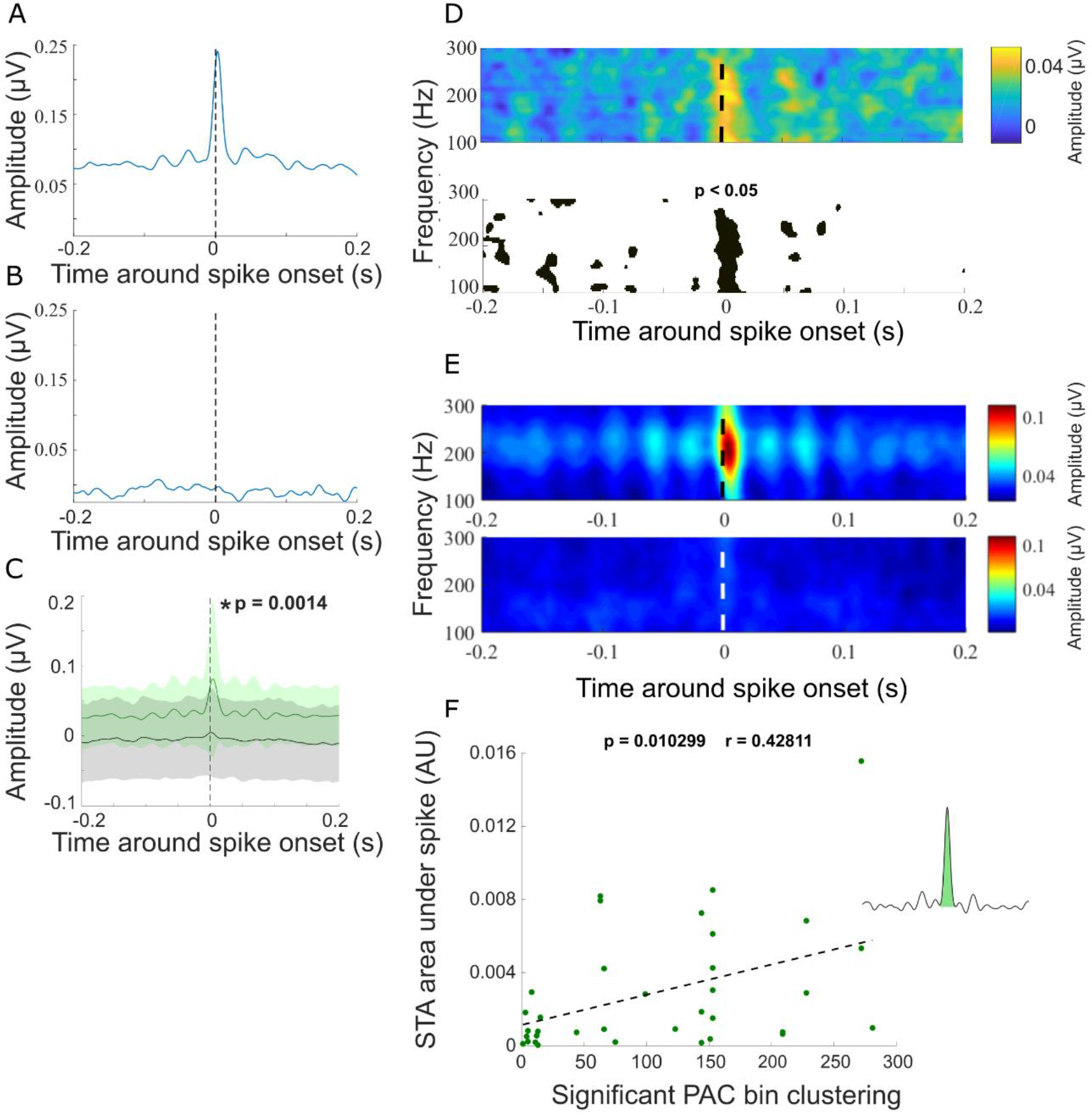
Spike-trigged HFO amplitude is increased during periods of PAC. **A-B)** Two example spike-triggered averages (STA) from recordings with (A) and without (B) PAC. The band pass filtered LFP HFO (100-300 Hz) amplitude was averaged around each single-unit spike, indicated by zero on the x-axis. **C)** Grand mean of all spike triggered HFO amplitude averages for recordings with significant PAC (green) +/− standard deviation (n = 39) and without significant PAC (black) +/− standard deviation (n = 35). * p = 0.0014 unpaired t-test at time zero. **D)** Example recording of spike triggered HFO amplitude (upper) and its significance testing (lower) across epochs with black on the colour scale indicating p < 0.05 **E)** Grand average of spike triggered amplitude across HFO frequencies from all recordings containing significant PAC (upper) and non-significant PAC-clustering (lower) with time zero denoting spike onset. Notice the repeating side peak-intervals, corresponding to beta frequency of ~30 Hz. **F)** The area under the spike of the spike triggered average significantly relates to the degree of significant PAC clustering calculated from microelectrode recordings (p = 0.0103, r = 0.4281, n=33).

In addition to averaging the HFO amplitude around individual spike-onsets, we extended the GLM-method to investigate how spike-amplitude coupling compared to GLM-calculated PAC patterns across all HFO frequencies and found a clear concordance between the significant spike-amplitude curves and significant PAC plots as exemplified in one recording from one patient in Figure 1, where significant PAC levels corresponded to spike-amplitude coupling peaks. This pattern was reproduced on the grand average level with significantly greater spike-amplitude coupling in the PAC group compared to the non-PAC group for HFO frequencies ranging from 120-224 Hz (t-test, p < 0.05 n(Non-PAC) = 38 n(PAC) = 39) (Fig. 3C), mimicking the PAC peaks for HFOs in the PAC comodulograms. Interestingly, spike-HFO-amplitude coupling peaked at the same 220 Hz frequency for which the maximal PAC values were found (Fig. 3 C-D). Taken together these findings suggest that, in the presence of PAC, HFO amplitude and the timing of single unit activity are comodulated by beta phase.

**Figure 3.**
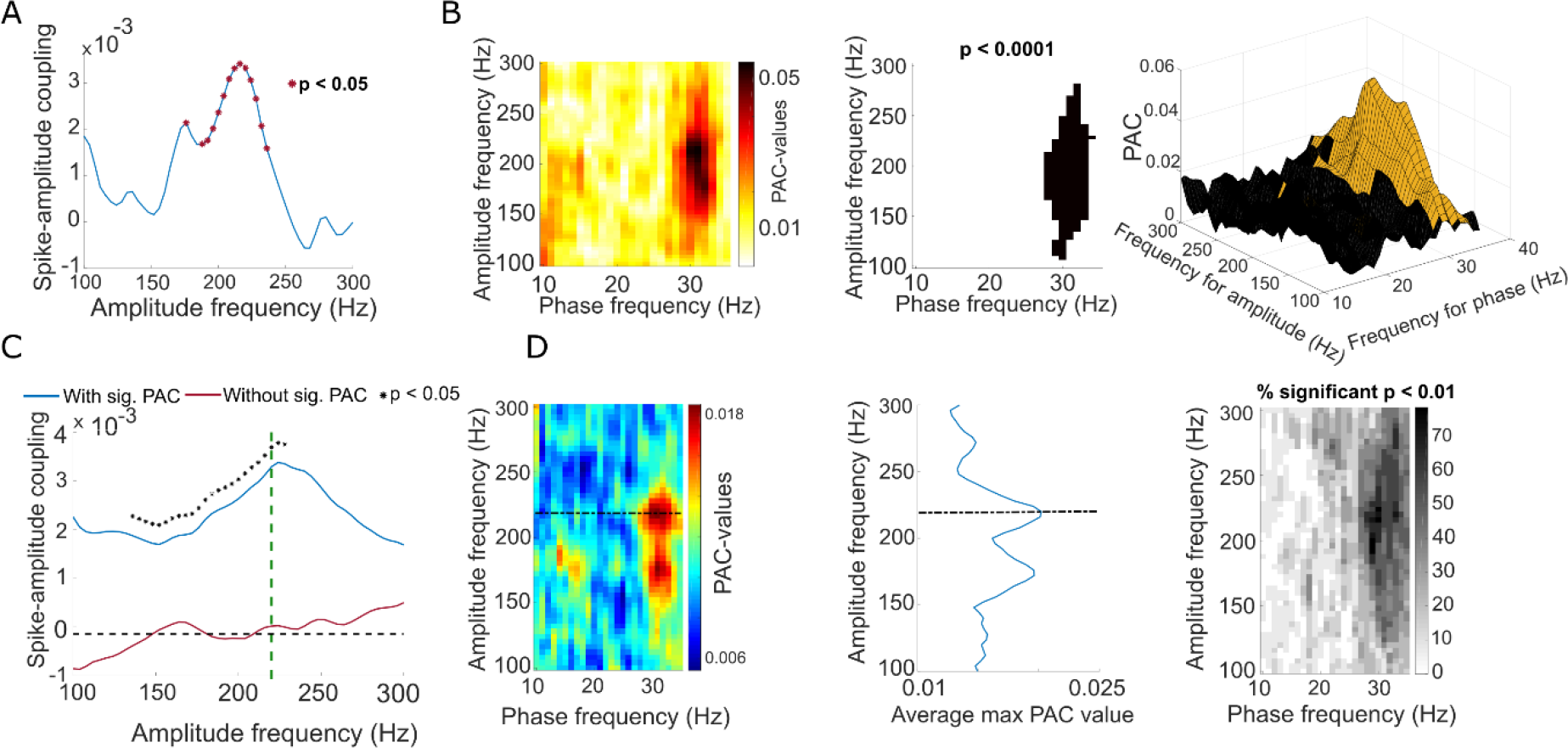
Contribution of spiking to PAC is revealed by multivariate analysis. Statistical coupling between single unit spiking and LFP HFO bandpass filtered amplitude - recorded at distance of at least 2 mm from each other - is seen to be significantly higher in the presence of significant PAC. Also, spike-amplitude coupling peaks at the same HFO frequency as maximum PAC-values. **A)** Example of spike-amplitude coupling plot across HFO frequencies.* denotes values that are significantly different from 0 (p < 0.05, t-test). **B)** The corresponding PAC comodulogram for A, PAC significance testing with p < 0.0001 (middle) and combined 3D PAC and p-value (brown: p < 0.01) representation. Note that spike-amplitude coupling peaks around the same HFO frequency (220 Hz) as the maximum PAC value in the comodulogram. **C)** Group averages of statistical spike-amplitude coupling for significant PAC (blue, n = 39) and non-significant PAC (red, n = 38) data sets (* p value < 0.05, two-sample t-test). The green dotted line marks the HFO frequency corresponding to the maximum PAC value from **D)** the grand averaged significant PAC comodulogram (left) with grey scaled frequency bins displaying percentages of significant p-values < 0.01 (right) and averaged max PAC-values across HFO-frequencies (middle).

Given that, in the presence of PAC, spike times predicted periods of high-amplitude HFOs lasting a few milliseconds, we hypothesised that increases in the number of spikes would lead to higher HFO amplitude over longer timescales. To this end, within individual recordings we assessed the relationship between the firing rate of STN neurons and HFO amplitude (180-220 Hz) in the neighbouring LFP (Fig. 4). Firing rates when PAC was present and absent were not significantly different (Average mean firing rate (PAC) = 29.15 Hz, non-PAC = 26.06 Hz, p = 0.5155, two-sided test). However, in recordings where PAC was present, we found an overall positive relationship between firing rate and HFO power across epochs of 3 seconds (PAC example Fig 4A, r = 0.6115, p < 0.0001, F-test). In the absence of significant PAC clustering, however, no relationship between overall between firing rate and HFO power could be found (Non-PAC example Fig. 4B, r = 0.0544, p = 0.6520). Across all recordings, we found r-values to be normally distributed around zero for the non-PAC group (Non-PAC mean-r = −0.0323, n = 62, Fig. 4C), but shifted to the right for recordings in which PAC was present (PAC mean r = 0.1906, n = 39, Fig. 4C). In line with this result, r-values in the presence of significant PAC were significantly greater than in its absence (p < 0.0001, two-sample t-test). Moreover, across recordings, we found that the span of significant PAC clustering and mean firing rate values were positively correlated for the PAC group (defined as clustering exceeding 30 frequency bins, p = 0.018, r = 0.408, n = 33, Fig. 5) but not the non-PAC group (defined as clustering below ten significant frequency bins, p = 0.789, r = 0.048, n = 31, Fig. 5).

**Figure 4.**
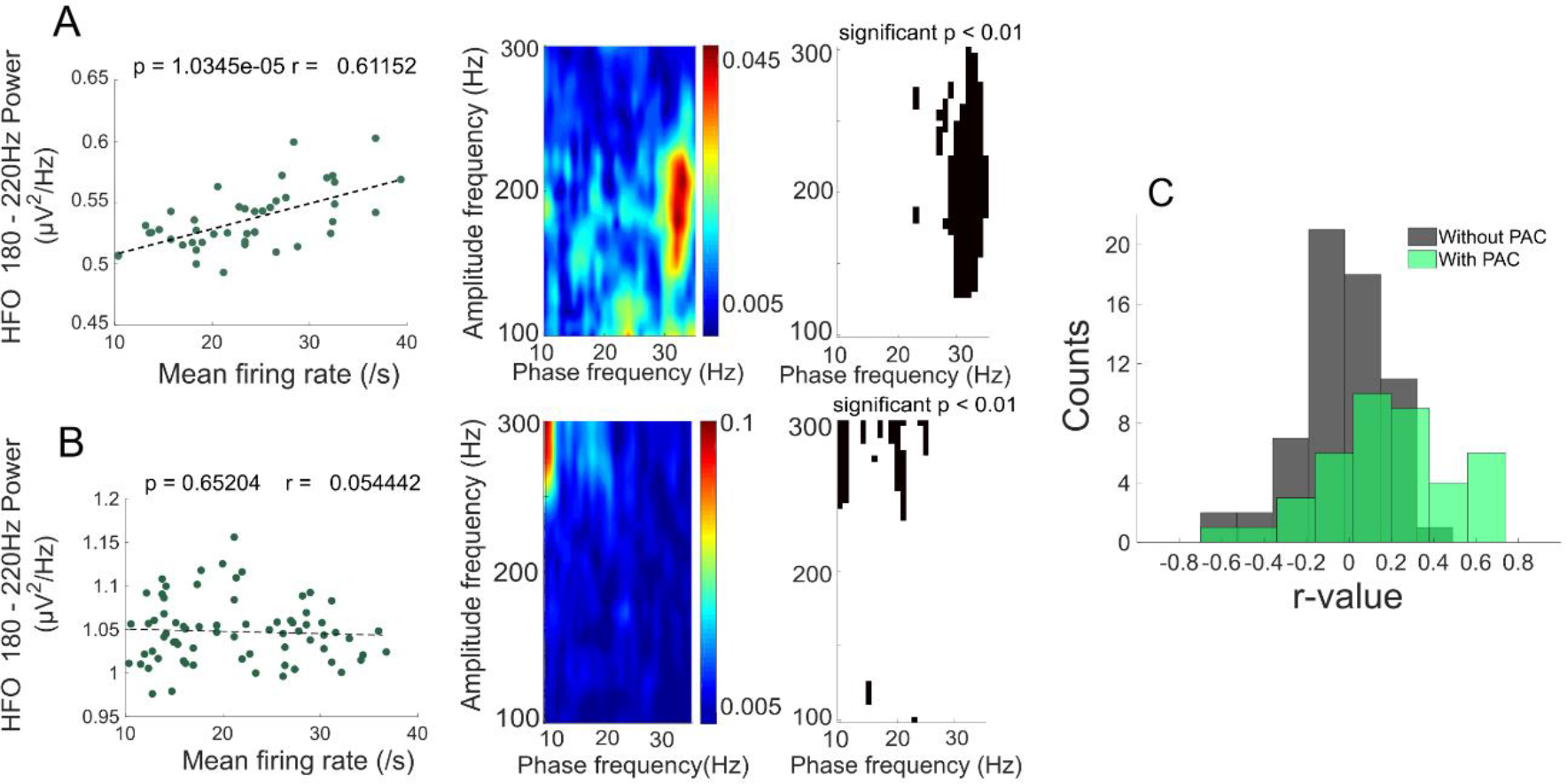
The time course of firing rate predicts HFO amplitude in the presence of PAC. A higher degree of synchronisation of STN spiking with high frequency oscillations is seen when the latter is modulated by the phase of beta frequencies. **A**) Example correlation between mean firing rate and HFO (180-220 Hz) amplitude correlation (p < 0.0001, r-value = 0.61152), each dot represents an epoch of 3 seconds. The corresponding PAC comodulogram and significance plot is on the right. **B**) As in A, but for a recording where PAC was absent. **C**) Population histogram of r-values from the correlation between mean firing rate and HFO power in the presence (green, n = 39, mean r = 0.1906) and absence (grey, n = 62, mean r = −0.0323) of significant PAC for all recordings. Note that r-values in the presence of significant PAC clustering are shifted to the right compared to the non-significant PAC data (p < 0.0001, two-sample t-test).

**Figure 5.**
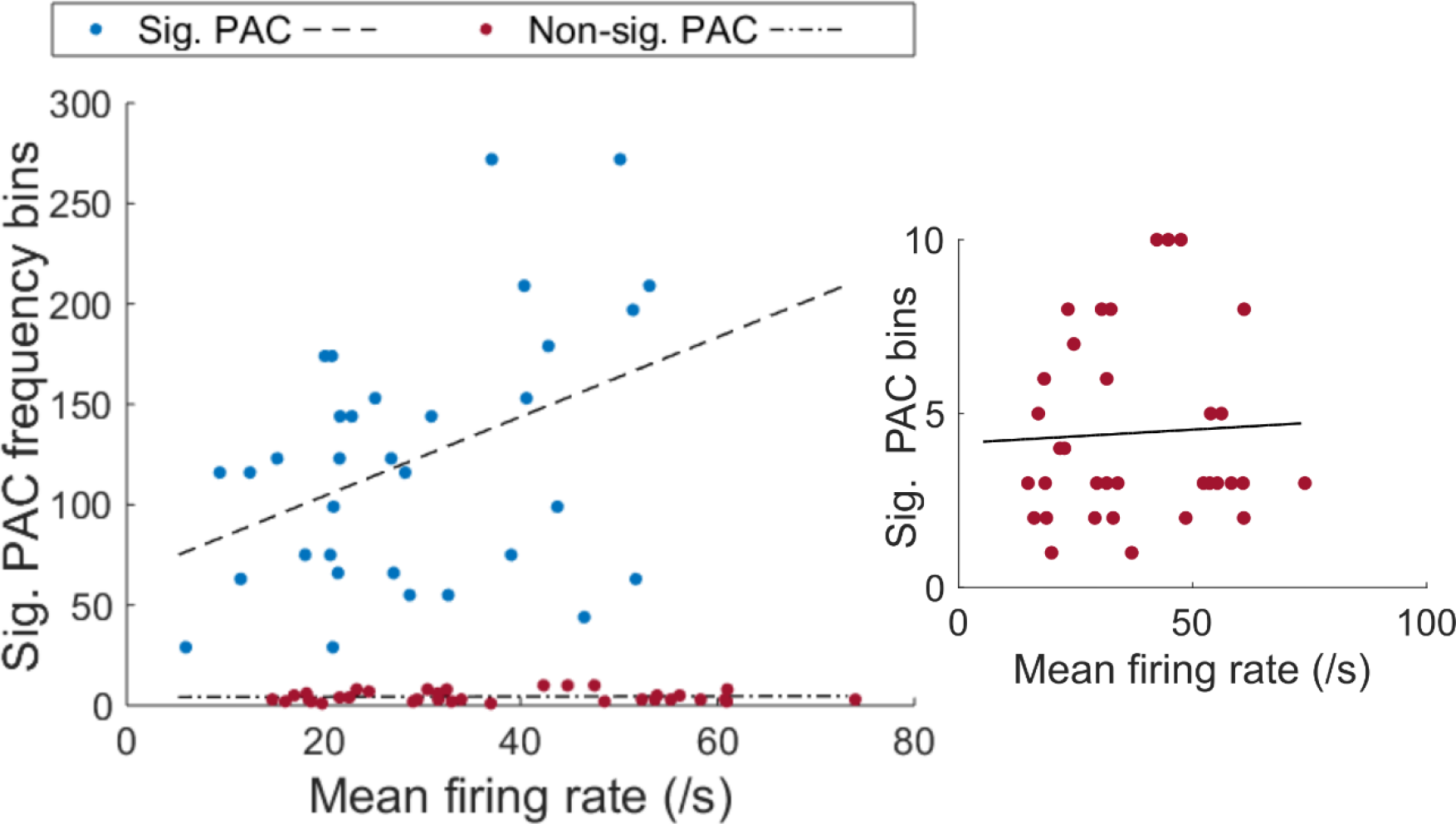
Extent of PAC predicts firing rate across whole recordings. Extent of significant PAC clustering correlates with mean firing rate when clustering exceeds 30 contiguous frequency bins (blue dots, p = 0.018, r = 0.408). In the case of fewer than 10 contiguous significant PAC bins (red dots, no significant PAC) there was no relationship between mean firing rate and the extent of PAC clustering (red dots, p = 0.789, r = 0.048). PAC and mean firing rate calculated from different electrodes.

Overall, increased spiking activity predicted the amplitude of HFOs only when PAC was present. These results suggest that spiking does not predict HFO amplitude *per se*, but only under conditions that also lead to significant PAC.

### Specific features of PAC are associated with the presence of synchronised beta oscillations in STN neurons

Next, we tested whether the phase and frequency parameters of PAC could be explained by their association with the spiking patterns of STN neurons. First, we examined whether there was a correlation between the angle of phase locking of the spikes and HFOs in recordings with and without PAC. In recordings with both PAC and phase locked units, both the spikes and HFOs were mostly clustered around the peak of the LFP beta oscillation (Fig. 6A) and the differences in their peak phases was not significant (p = 0.1889, Watson-Williams test). In contrast, in recordings where no PAC or phase locking was present (Fig 6B-D), both units and HFOs were locked to all phases except for this peak and the differences in preferred phase were significant (p = 0.0124, Watson-Williams test) (Fig. 6B). This analysis further indicates that STN spiking underlies the time course of HFO amplitude under conditions where STN spiking is locked to local beta oscillations.

**Figure 6.**
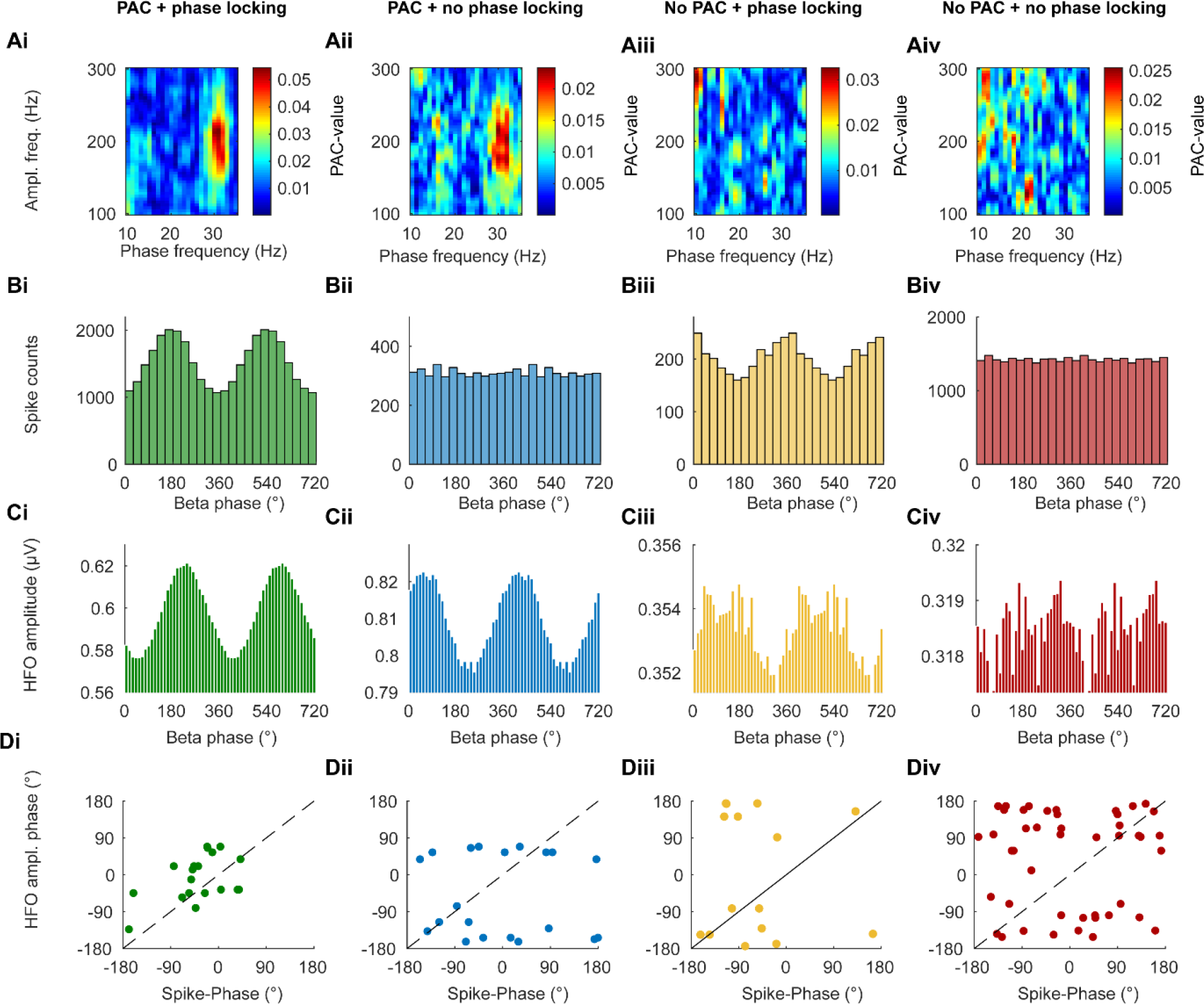
Phase-locked spiking and high HFO amplitude cluster around the same phase of the LFP beta oscillation. The figure is divided into columns with regards to whether PAC and/or spike-phase locking were significant in a pair of LFP/unit recordings. **i)** Significant PAC and significant spike phase-locking, column **ii)** significant PAC without significant phase-spike locking, column **iii)** no significant PAC but significant phase-spike-locking and column **iv)** no significant PAC and no significant phase locking. **A)** PAC comodulogram examples. **B)** Spike-phase-plots for same recordings as in A. **C)** Beta-phase HFO-amplitude histograms for the same recordings. **Di-iv)** Mean phase of spiking plotted against the phase with the highest HFO amplitude for each recording meeting the criteria labelled at the top of the column. Diagonals mark complete phase alignments. Note that Di shows that both spikes and the highest HFO amplitude occur around 0° when both PAC and spike-phase locking are present.

A consistent feature of STN LFP PAC is that the HFO frequency peaks > 200Hz (Lopez-Azcarate et al., 2010; Özkurt et al., 2011; Yang et al., 2014; van Wijk et al., 2016), considerably higher than in cortex (Manning et al., 2009; de Hemptinne et al., 2015). Having established a temporal correlation between beta-locked STN spikes and HFOs, we next investigated whether the spectral properties of these phase-locked neurons could explain the frequency of STN HFOs. To this end we compared the power spectra across single unit recordings in recordings with and without significant PAC (Fig. 7). The single units at sites with PAC showed both a clear peak between 150 and 250Hz and a peak at beta frequency (Fig. 7A). Both peaks were significantly greater than in the non-PAC group (p < 0.05, F-test, Fig. 7B), which did not show an obvious peak above 100Hz. In recordings with PAC, the peak frequency of the unit power spectra was significantly correlated with that of the HFO (Fig. 7C), while there was no such correlation when PAC was absent.

**Figure 7.**
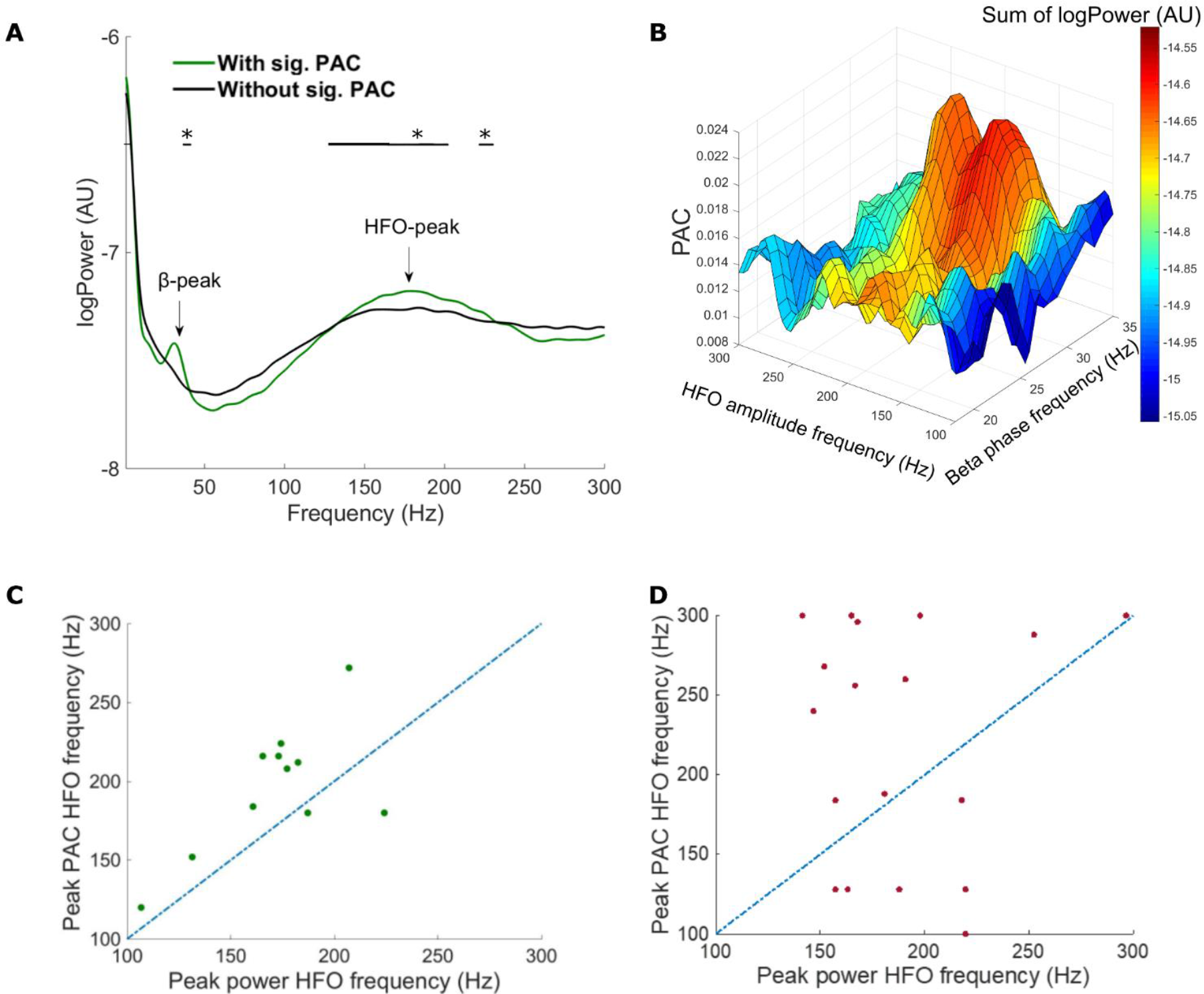
The power spectral density features of single unit activity predict HFO frequency. **A**) Group averaged single unit spectral densities at PAC (green) or non-PAC (black) clustering recorded sites. Note that only single units at PAC sites demonstrate a beta peak and a larger gamma peak. * p < 0.05, Two-sample F test for equal variances. **B**) 4D representation of grand average microelectrode phase-amplitude coupling for the significant PAC group and the overlapping power spectrum summation with x-values = beta-phase frequency, y-values = HFO amplitude frequency, z-values = PAC-values. The single unit spectral power densities for the beta and HFO frequencies represented by the colour bar (c-axis). The graph shows how the PSD intensity corresponds with PAC-values. **C + D**) Peak PAC HFO frequencies plotted against peak power HFO frequencies for microelectrode recordings from the significant PAC group (C, n = 11) and the non-significant PAC group (D, n = 18) demonstrating a higher degree of correspondence between HFO power peak frequencies and PAC peak HFO frequencies for the significant PAC-data. Proximity of dots to the blue line indicates a correspondence between peaks in PAC and power.

### Increased PAC is predicted by the duration of beta bursts

Recent studies have demonstrated that beta oscillations are not continuous, but occur in transient, high amplitude bursts of several cycles that are thought to reflect transient synchronisation of synaptic inputs to the STN (Tinkhauser *et al.*, 2017a; Tinkhauser *et al.*, 2017b). We next asked whether the occurrence of PAC was independent to that of beta bursts, or whether the two measures could reflect the same underlying process. Here, we calculated PAC by the PLV-method, which is preferable for short time series. For individual recordings, calculation of PAC inside and outside beta bursts in the same recording led to clearly different comodulograms, with considerably larger PAC in the burst periods (Fig. 8A). This result was consistent across the data set, with significantly higher PAC during beta-bursts than outside bursts across hemispheres in the presence of a beta-peak (PAC during bursts = 0.2055, PAC outside bursts = 0.0989, p = 0.0234, paired t-test) (Fig. 8B).

**Figure 8.**
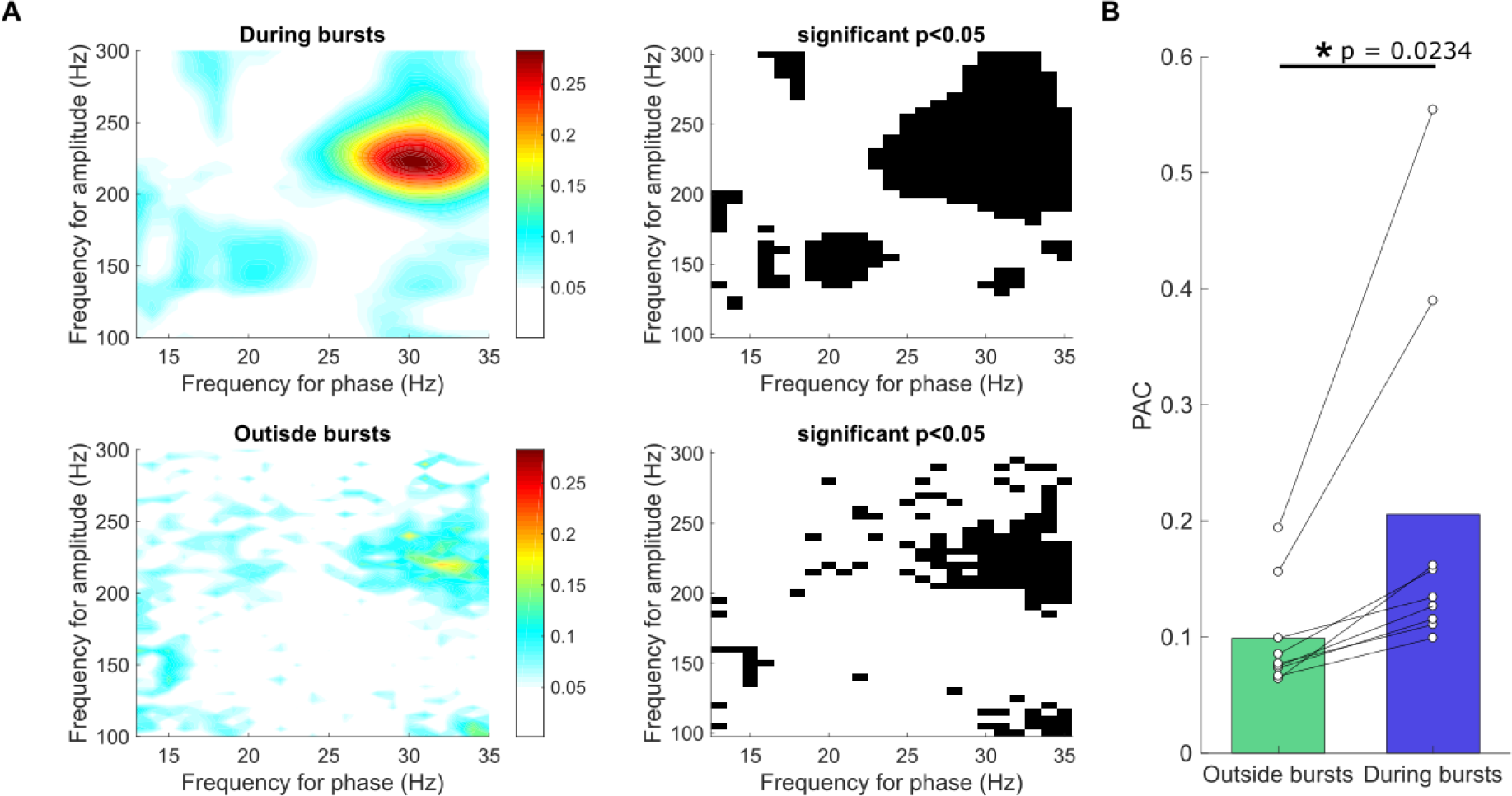
PAC is significantly higher during beta bursts. **A**) Example PAC comodulogram inside and outside bursting period from the same recordings, calculated by the PLV method. B) Average around maximum (± 16 Hz) PAC calcuated from 9 hemispheres with recordings that demonstrated a beta peak. PAC was found to be significantly higher during periods of beta bursting compared to outside periods of bursting (PAC during bursts = 0.2055, PAC outside bursts = 0.0989 p = 0.0234, paired t-test, n = 9) in the presence of a beta peak.

Moreover, we found that there was a significant increase in PAC with burst duration and when progressively higher thresholds were used to define the burst epochs, following a correction for the shortening of the data segments as the threshold and durations were increased (One-way ANOVA, p < 0.0001, Fig. 9). Longer bursts and higher amplitude thresholds consequently captured higher levels of synchronisation between the beta phase and amplitude of the HFOs. Finally, we also found progressive increases in both HFO and beta power with increasing burst durations (Fig. 10). Overall correlations between different measures in the paper are summarised in Table 1. In summary, PAC was higher within beta bursts and increased progressively with the magnitude (set by threshold) and duration of the bursts.

**Figure 9.**
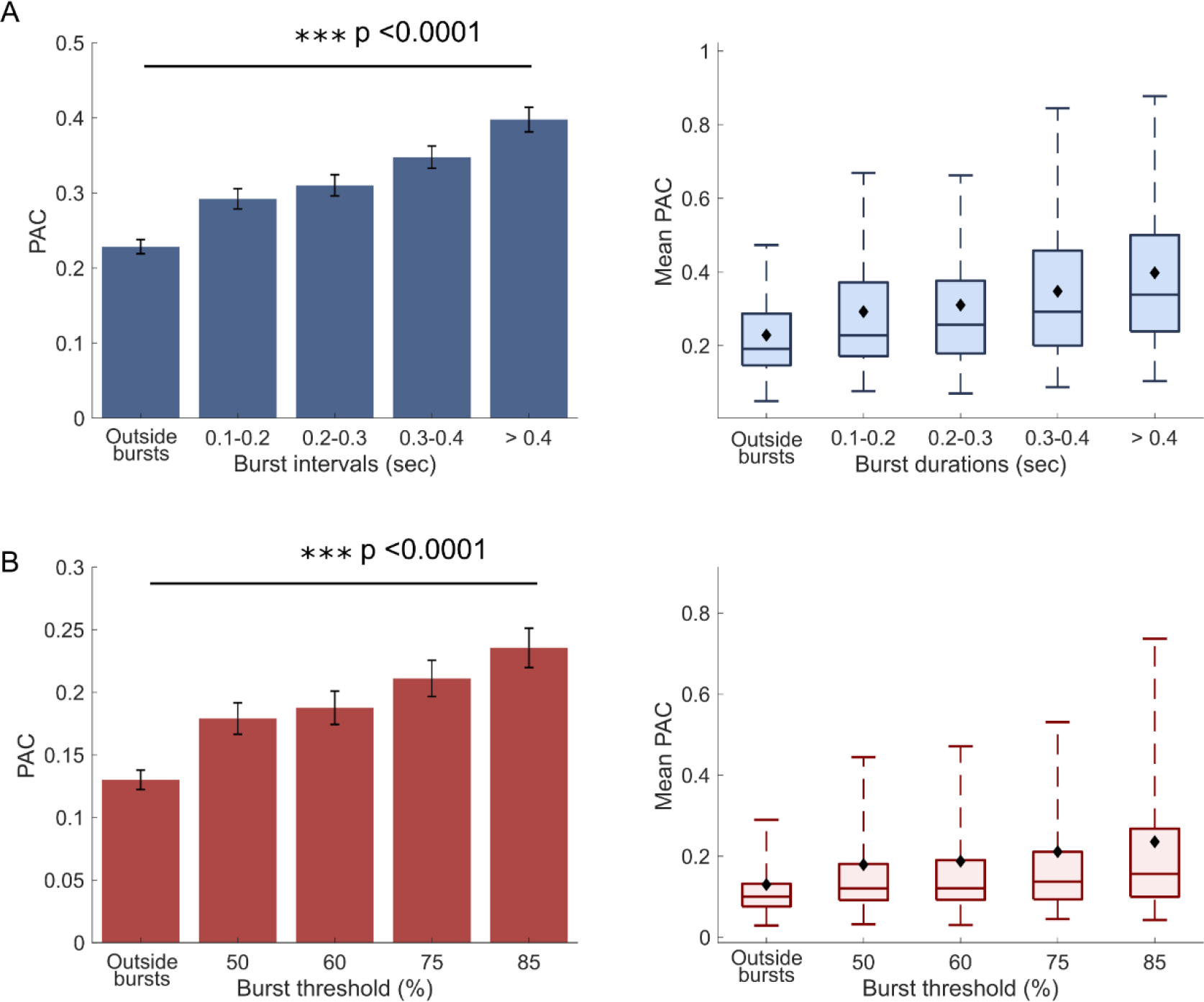
PAC increases progressively with both burst duration and thresholds increments. **A,** Bar and box plots of PAC at different burst durations. PAC significantly increases with burst duration with longer bursts capturing higher levels of coupling. **B**, Bar and box plots of PAC at different burst thresholds. PAC increases significantly with threshold increments. All concentrated burst vectors were length adjusted.

**Figure 10.**
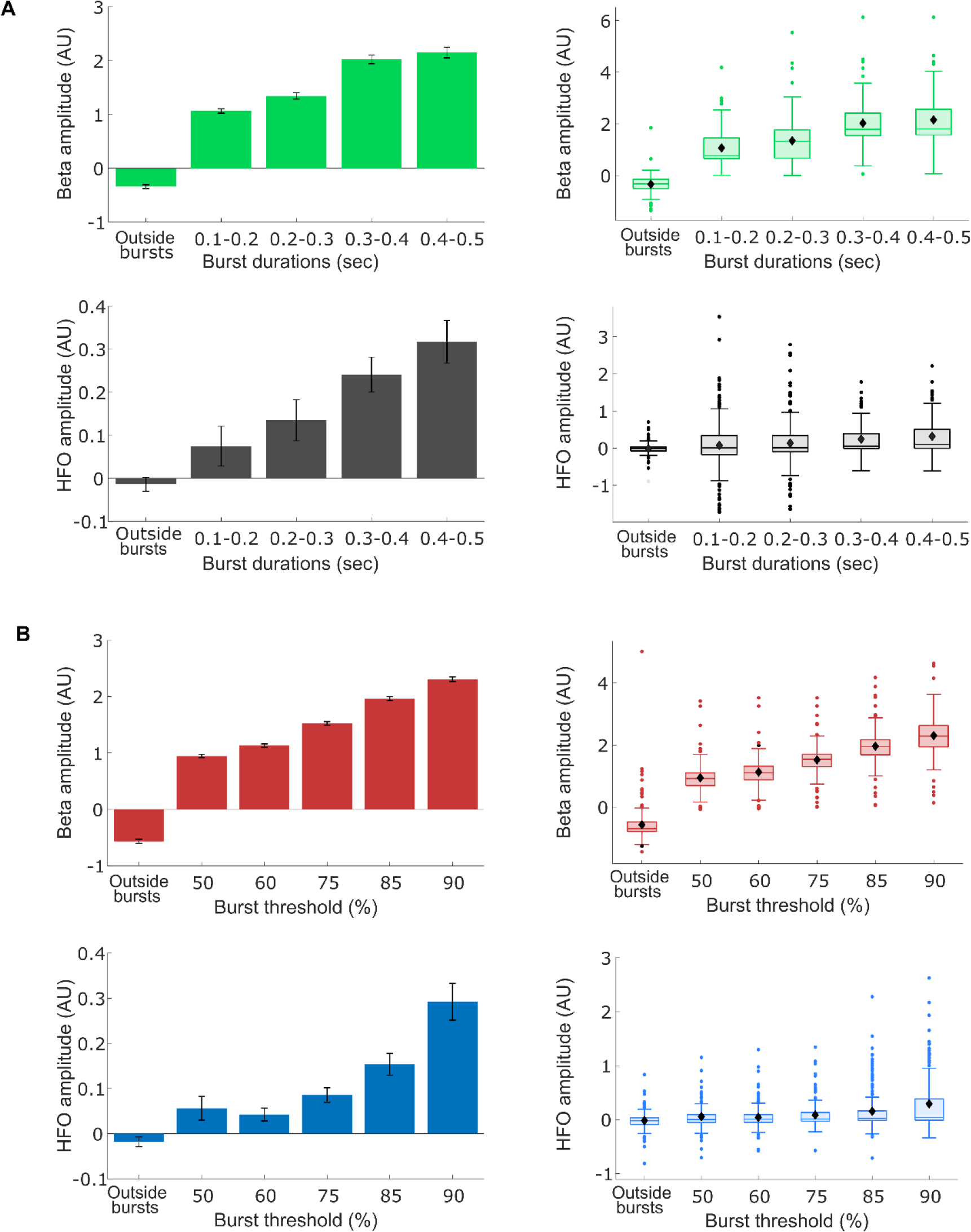
Both normalised beta and HFO amplitude increase significantly with both burst duration and thresholds. A) Bar and box plots of average beta (green) and HFO (grey) amplitudes for different burst durations. B) Bar and box plots for average beta (red) and HFO (blue) amplitude for different burst thresholds.

**Table 1:** Associations between measures of PAC, spiking activity and beta bursts. Ticks and crosses show when two measures, denoted by the row and column labels, were/were not associated based on analysis of PAC vs non-PAC recordings or temporal/phase/frequency correlations within recordings.

|  | Spiking Activity |  | Spectra/Phase |  | Beta Bursts |  |  |
| --- | --- | --- | --- | --- | --- | --- | --- |
| **PAC Parameters** | Spike Times | Mean Firing Rate | Spike Phase | Unit high frequency peak | Beta-burst In/out | Beta-burst Amp. | Beta-burst Length. |
| PAC | N/A | ✓ | ✓ | N/A | ✓ | ✓ | ✗ |
| HFO Amplitude | ✓ | ✓ | N/A | N/A | ✓ | ✓ | ✓ |
| HFO Frequency | N/A | N/A | N/A | ✓ | N/A | N/A | N/A |
| HFO Beta Phase | N/A | N/A | ✓ | N/A | N/A | N/A | N/A |

## Discussion

In this study, we investigated how different aspects of PAC in the STN LFP related to the spiking activity of STN neurons. We analysed the relationship between STN beta oscillations, HFOs and spiking activity in the presence and absence of PAC and investigated how PAC differed inside and outside of beta bursts. Our results suggest that PAC reflects the oscillatory recruitment of STN spiking activity at the beta frequency and that this coupling is stronger during transient beta-burst events. These findings raise specific issues regarding the relationship of PAC to other biomarkers of pathophysiological activity in PD.

### Physiological basis of PAC in the STN

PAC has been demonstrated in many brain areas including several cortical regions (Canolty *et al.*, 2006; Canolty and Knight, 2010; Szczepanski *et al.*, 2014; Voytek *et al.*, 2015), basal ganglia (Tort *et al.*, 2008; von Nicolai *et al.*, 2014) and hippocampus (Tort *et al.*, 2008; Scheffer-Teixeira *et al.*, 2012; Colgin, 2015). Across this range of structures, there is considerable variability in the frequency of both the phase and amplitude signals. Several biological and nonbiological mechanisms can lead to significant PAC in a given time series. The first common interpretation is that there is functional coupling between neural populations oscillating at different frequencies. Such coupling has been convincingly demonstrated in the hippocampus, where the power of gamma frequencies is locked to specific phases of the ongoing theta oscillation, depending on the location of the recording electrodes (Buzsaki and Wang, 2012; Scheffer-Teixeira *et al.*, 2012; Colgin, 2015). In this case, both carrier and amplitude oscillation are underpinned by the synchronised firing of the underlying neurons. A second common interpretation is that higher frequency gamma oscillations are generated by background spiking (Ray *et al.*, 2008a; Ray *et al.*, 2008b; Ray and Maunsell, 2011) and PAC is the result of the locking of this activity to a low frequency carrier oscillation. Nevertheless, here the two oscillations represent semi-independent processes, i.e. the carrier oscillation is dominated by synaptic input and the high frequency oscillation by background spiking. The third interpretation is that the shape of the carrier signal creates a high frequency spectral component that has a consistent phase (Aru et al, 2015). This was recently demonstrated for cortical PAC in PD patients, where the asymmetry of the beta oscillation was shown to create the phase locked high frequency spectral content (Cole et al., 2017; Wijk, 2017).

In regard to STN PAC, we provide several lines of evidence for the second interpretation; that beta oscillations, representing coordinated synaptic input, drive spiking activity, which manifests as high frequency oscillations. Firstly, only when PAC was present was spiking correlated with HFO amplitude. It is worth reiterating that the spike was at least 2 mm away from the LFP channel, eliminating the possibility of direct electrical contamination. Rather, the time of spikes at a distant site predicted the timing of high HFO power with millisecond precision. Importantly, peaks were not seen when PAC was absent, suggesting that spiking only leads to peaks in HFO in certain contexts. Secondly, over time windows of a few seconds, firing rate was positively correlated with HFO power in the presence of PAC. As these correlations were absent when PAC was not present, such correlations are more likely the result of correlated, rather than any spiking, which is caused by beta synchronisation. Moreover, there were no significant differences in mean firing rate between the PAC and non-PAC groups, highlighting that this result must be due to temporal dynamics, rather than excitability per se. These conclusions are supported by the finding that the phase of HFO locking to the beta carrier predicts that of single units. Finally, we show that the spiking of single STN neurons can produce spectral frequencies that match those of the HFO. This is a novel and important finding, as the HFO frequency is much higher in STN than in cortex. It may reflect the high frequency firing of STN neurons when they are recruited to beta, which correlates with motor symptom severity (Sharott et al., 2014). The correlation between bursting and PAC is supported by a recent computational study demonstrating that spike bursts with the intraburst and interburst intervals seen in STN neurons in PD patients can generate PAC (Sanders, 2016).

Our findings are different to those of Yang and colleagues (Yang et al., 2014) who suggest, using similar data to our own, that spike locking to beta oscillations is independent from HFO locking. There could be several reasons for this discrepancy. Here we only used LFPs that were recorded within the STN, as defined by the characteristic spiking of STN neurons. Although we concur with Yang and colleagues that the LFP dorsal to the STN probably contains volume conducted activity from that structure, it may likely have a greater contribution from other sources such as thalamus and zona incerta. The GLM method and the commonly used modulation index (as determined by the circular mean method), are in essence the same when the latter is properly normalised, and similar results have been yielded by both methods (Penny et al., 2008). However, we used a more stringent criterion for defining significant PAC, which may have reduced our calculations of the HFO beta phase to those with the strongest locking, and thus most stable phase. Given the larger data set in Yang et al, further characterisation of STN PAC, possibly in animal models where more single neurons can be recorded, would be highly valuable for resolving this issue.

### PAC during episodes of transient input synchronisation

Synchronisation across STN synaptic inputs is thought to be indexed by beta LFP power (Weinberger *et al.*, 2006; Mallet *et al.*, 2008a; Mallet *et al.*, 2008b; Tinkhauser *et al.*, 2017b; Sharott *et al.*, 2018). Recently, much attention has been focussed on the interpretation of neuronal oscillations as transient, burst-like events (van Ede et al., 2018). This has been found to be particularly true of oscillations in the beta range, both in the healthy (Feingold *et al.*, 2015; Mirzaei *et al.*, 2017) and Parkinsonian (Tinkhauser *et al.*, 2017a; Tinkhauser *et al.*, 2017b; Tinkhauser *et al.*, 2018) cortico-basal ganglia network. The transient nature of these oscillatory events, suggests that these bursts are periods where synaptic inputs are most synchronised and that it is these sustained periods of synchronisation in PD that impairs movement processing. We found that beta bursts also have significantly higher PAC and that the PAC increases progressively with duration and amplitude (assessed by increments in burst threshold) of the bursts. In line with this, dopamine replacement in PD patients supresses both burst length (Cagnan *et al.*, 2015; Tinkhauser *et al.*, 2017b) and PAC (Lopez-Azcarate et al., 2010; van Wijk et al., 2016). Overall, PAC and beta bursts may thus reflect periods of increased input/output synchronisation, or transfer, in the beta frequency range. Tinkhauser and colleagues showed that that adaptive DBS truncated longer bursts in favour of bursts of shorter durations and lower amplitudes (Tinkhauser et al., 2017a). As longer bursts proportionately capture higher levels of input/output synchronisation, as indexed by PAC, our analysis suggests that successful stimulation strategies will reduce burst length and PAC magnitude in parallel.

### PAC as a biomarker of pathophysiology

PAC is increasingly suggested as a primary biomarker for pathophysiology in PD patients. At the level of STN, our results suggest that this may be due to PAC being a proxy for the locking of local spiking activity to beta oscillations, which is supported by computational evidence (Sanders, 2016). The importance of this interpretation is that beta oscillations, as measured in basal ganglia LFPs or EEG/ECoG, are synchronised across the entire cortical basal ganglia network (Mallet *et al.*, 2008a; Mallet *et al.*, 2008b; Brazhnik *et al.*, 2016; Sharott *et al.*, 2017) and such spike-field locking is a robust marker of the Parkinsonian state across experimental animals and patients. Using the timing of cortical spiking activity locked to such network oscillations as a biomarker for closed-loop deep brain stimulation relieves motor impairment to a greater degree than conventional 130Hz stimulation (Rosin *et al.*, 2011). PAC may thus serve as a useful way of detecting the locking of local spiking activity to network oscillations from wideband signals. In terms of its use as a biomarker for closed-loop stimulation, the main question is whether the HFO locking provides information that cannot be extracted from the beta oscillation alone, using metrics of amplitude, phase and possible cycle asymmetry. The possible masking of significant PAC by stimulation may also impede its use to control stimulation timing (Sanders, 2016). Nevertheless, given the proliferation of devices allowing long term recording of STN LFPs, our results suggest that PAC and beta-bursts can provide a useful parameter for measuring the synchronisation and/or oscillation of underlying unit activity in the absence of direct access to measurements of spiking.

## Acknowledgements

A. M’s DPhil-studies are funded by the Rosetrees Trust. A. S. and P.B. by the Medical Research Council (MC_UU_12024/1) and National Institute for Health Research Oxford Biomedical Research Centre. G.T. received a grant from the Swiss Parkinson Association. BW received funding from the European Union’s Horizon 2020 research and innovation programme under the Marie Sklodowska-Curie grant agreement No 795866. A.K.E. and C.K.E.M acknowledge a grant from the German Research Council (SFB 936, projects A2/A3, and C8 respectively).

---

**Supplementary figure 1.**
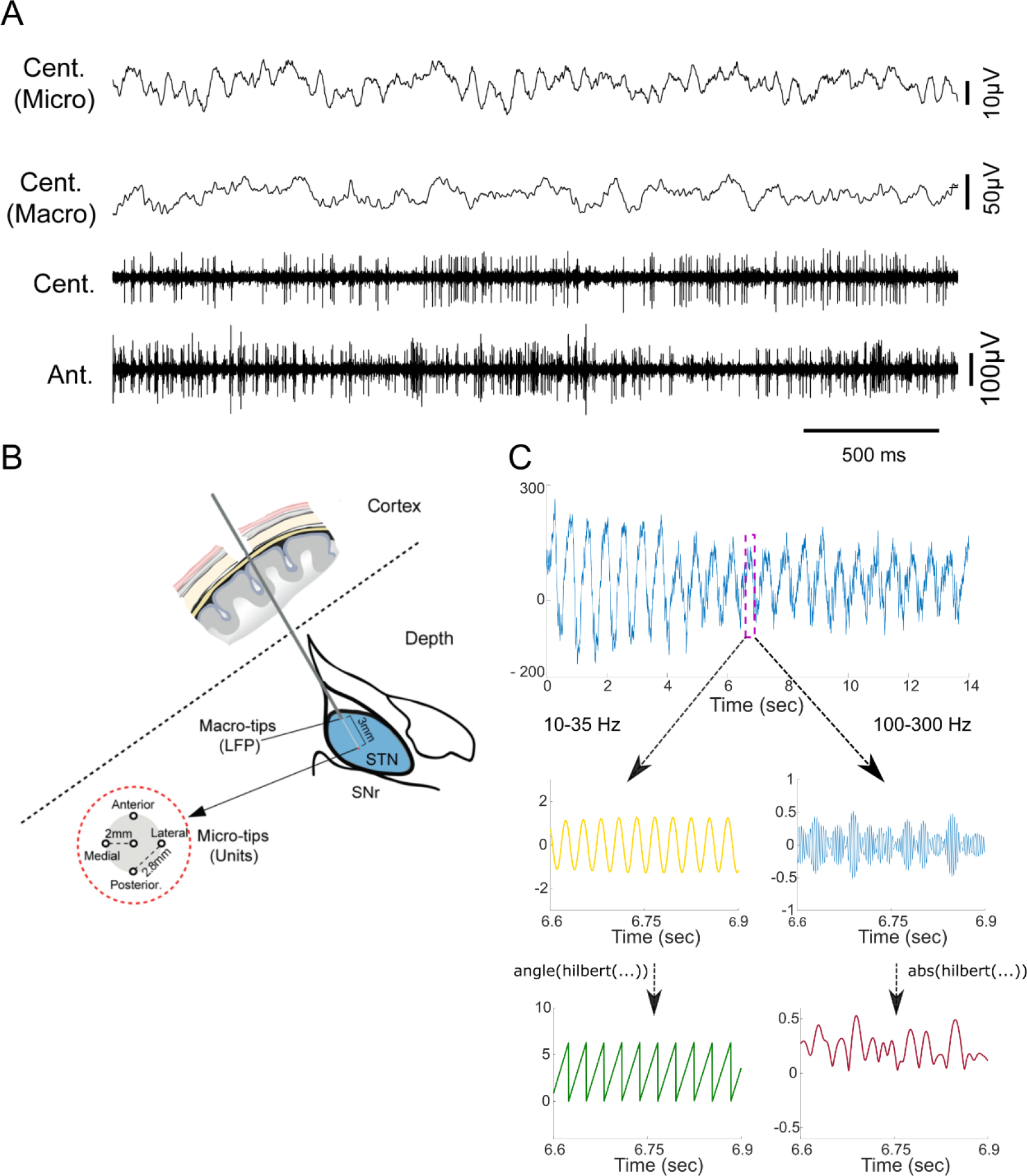
Local field potentials (LFPs) were recorded from the subthalamic nucleus (STN) from micro− and macro-electrode recordings during deep brain stimulation surgery. **A**) Example of micro- and macro electrode recordings of raw LFP signals in one patient and microelectrode multi- and single unit recordings from an anterior and central microelectrode, respectively. The micro electrodes were organised in a concentric array **(B)**with four outer micro electrodes distanced 2 mm from the central micro-electrode. All simultaneous spiking and LFP analyses were made from microelectrodes at least 2 mm apart. **C)** In order to apply the GLM-method to calculate PAC the LFP signal was bandpass filtered and Hilbert transformed in order to extract the instantaneous HFO (100-300 Hz) amplitude and broad beta phase (10-35 Hz).

**Supplementary Figure 2.**
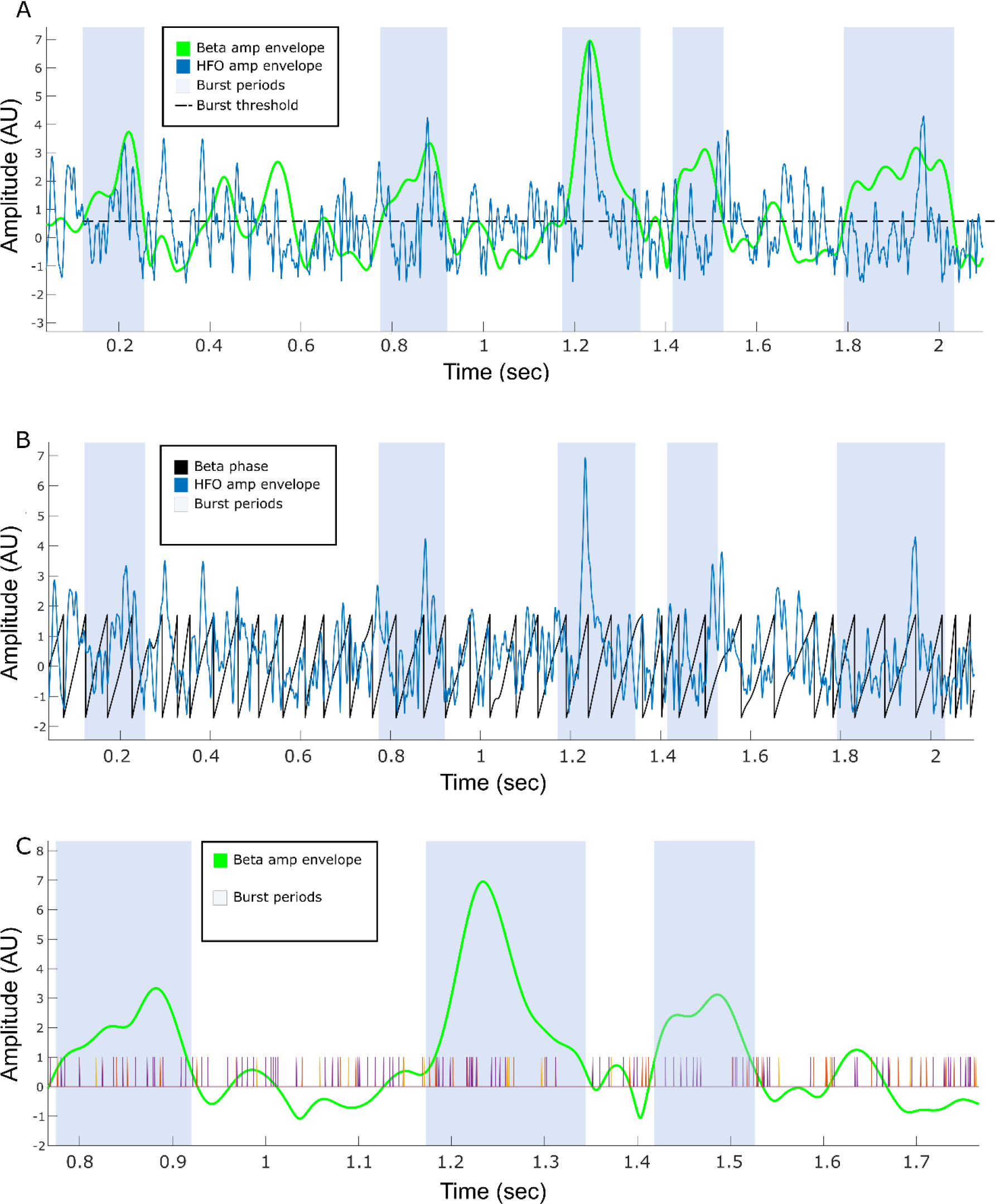
Example of occurrence of HFOs, spiking and beta bursts in single recordings. A) Example recording of broadband HFO (100-300 Hz) amplitude time series and HFO amplitude filtered around the beta peak frequency. Shaded blue areas mark periods of beta-bursting of more than 100 ms. The dashed line outlines the 75% burst threshold. B) Example of normalised HFO filtered amplitude envelope and beta phase. C) Beta amplitude envelope (green), beta bursts (shaded blue areas) and unit firing (orange, yellow and purple).

**Supplemental Table 1.**
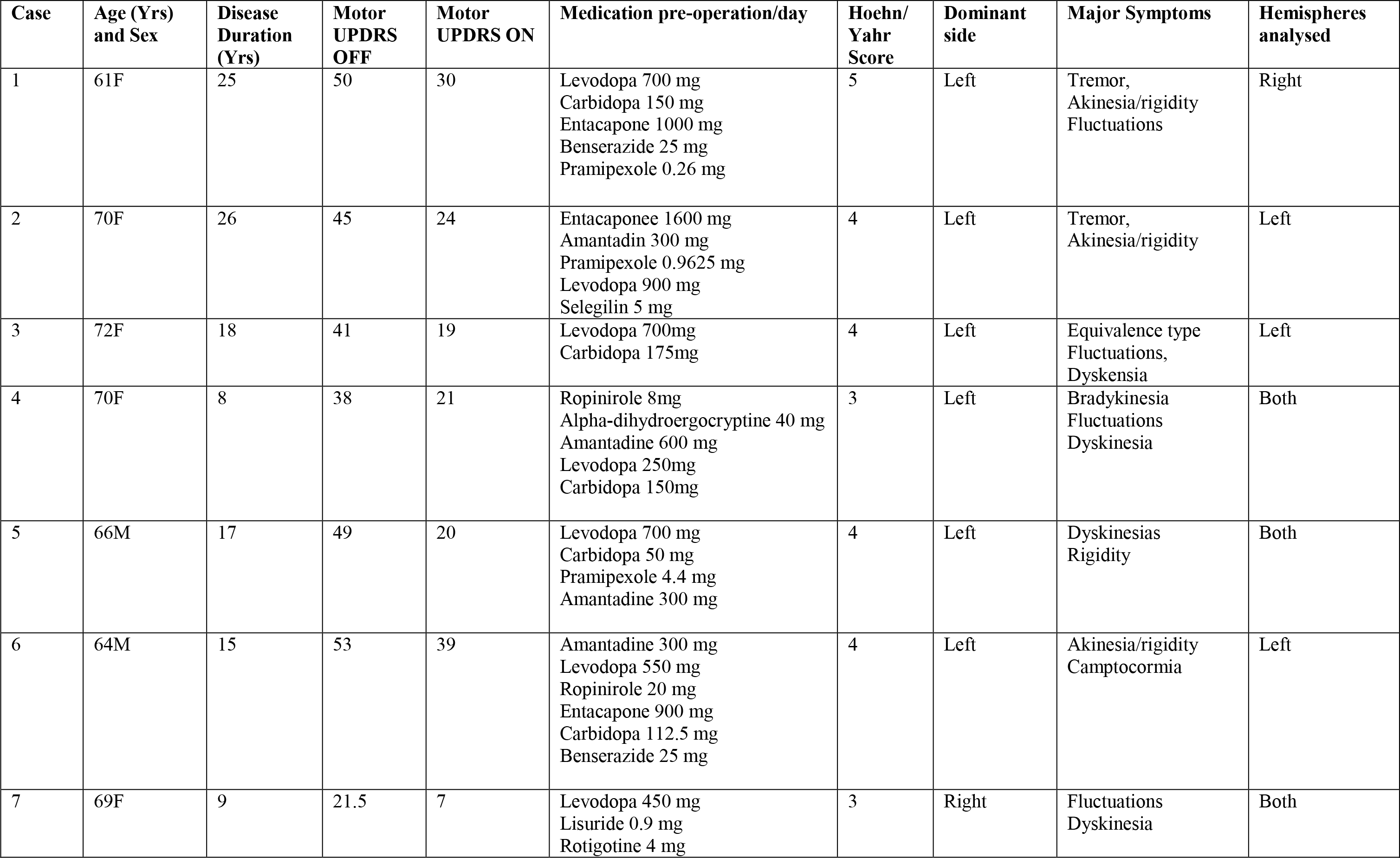
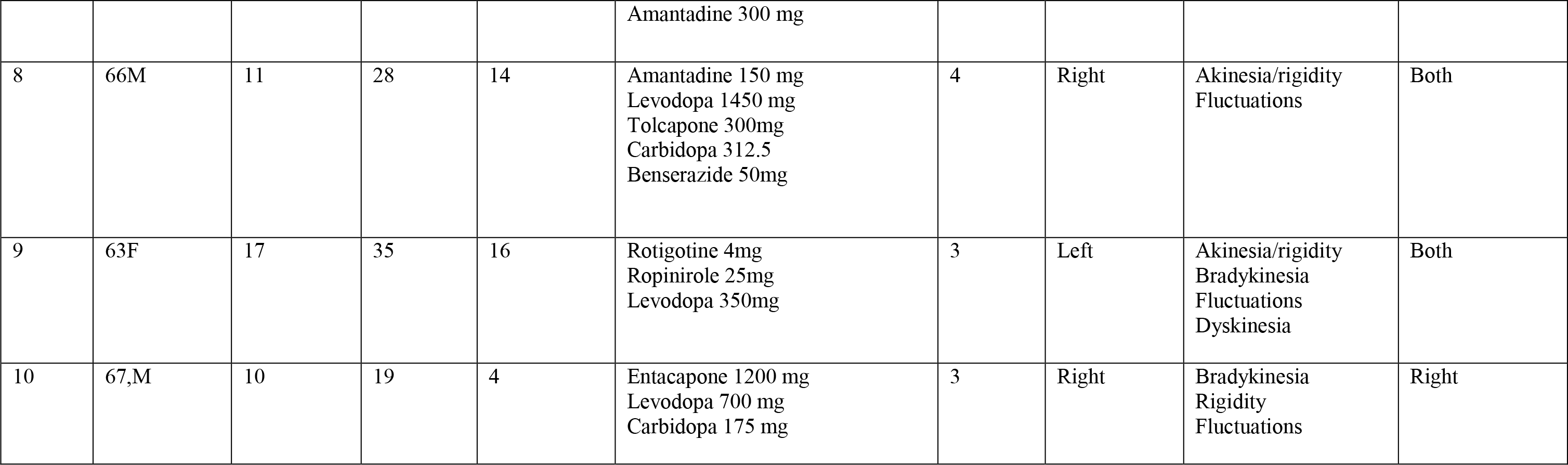
Patient details.

